# Modeling the dietary effects on bat viral shedding and potential consequences for pathogen spillover

**DOI:** 10.1101/2024.06.19.599703

**Authors:** Chiara Vanalli, Caylee Falvo, Dan Crowley, Benjamin Schwarz, Raina Plowright, Peter J. Hudson, Agnieszka Rynda-Apple, Isabella M. Cattadori

**Affiliations:** Center for Infectious Disease Dynamics and Department of Biology, The Pennsylvania State University, University Park, 16802 PA, USA; Department of Public and Ecosystem Health, College of Veterinary Medicine, Cornell University, Ithaca NY 14853 USA; Research and Technologies Branch, National Institute of Allergy and Infectious Diseases, National Institutes of Health, Hamilton MT 59840 USA; Department of Microbiology & Cell Biology, Montana State University, Bozeman MT 59715 USA

**Keywords:** Influenza virus, Immunity, Citrulline, Food consumption, Laboratory experiment, Data fitting

## Abstract

Changes in the quality and quantity of food resources can affect individuals’ health, the way they control infections and consequently the likelihood of onward transmission of pathogens. Dietary shifts have been proposed as one of the factors driving spillovers of zoonotic viruses from bats through a bridging host to humans. While there is a general understanding of the relationship between nutrition and infection in model systems, how diet affects pathogen shedding and the risk of spillover from bats is lacking. We used a data-driven mathematical modeling approach to disentangle the relation between diet, immunity, and viral shedding of Jamaican fruit bats infected with H18N11 and fed different dietary regimes. Model selection indicates that the synergistic interaction between the metabolite citrulline and the cytokine TNFα controls viral shedding in a diet-dependent manner. Bats on a sub-optimal fat diet are more successful in terminating shedding than bats on an optimal or sub-optimal sugar diet. However, when bat foraging behavior is considered, bats on the optimal diet show a lower spillover hazard, probably because of a feeding behavior less conducive to transmission. This study provides novel insights into the diet-driven mechanisms of viral shedding and how they can potentially contribute to spillover events.

## Introduction

Zoonotic spillovers, whereby pathogens are transmitted from an animal reservoir to humans, have been recognized as one of the primary causes for the emergence and spread of infectious diseases, and have been linked to recent outbreaks including SARS-CoV-2, Nipah, Hendra, MERS and H1N1-2009 (1–4). Bats are the reservoir hosts of several of these viruses, which cause severe disease in humans and great risk to public health (5,6). Successful spillover requires the pathogen to overcome a series of barriers, from the environmental factors that limit pathogen circulation in the animal reservoir (e.g. population density and host permissiveness), to the impediments encountered during the invasion and onward transmission in the human population (e.g. immune response and molecular compatibility) (1). Among these, is the fundamental condition of viral shedding from the animal reservoir, whereby a sufficient infectious dose results in an amplifying infection within the human host or a bridging host. Hypothetical modeling scenarios suggest that the greater the dose of viral shed over a time frame, whether this is a constant or a pulse process, the higher is the probability of cross-species transmission (1,7). If this dose-response relationship is linear, the probability of spillover success will be similar under constant or pulse shedding, while a non-linear relationship will result in a successful event if the dose is above a threshold, which might not be the same for constant and pulse shedding (1). Therefore, in addition to the processes that affect the degree and duration of shedding by an animal reservoir it is important to consider the conditions that produce a successful dose.

Shifts in the dietary requirements and/or access to adequate nutritional resources are frequently proposed to affect the way animals control infections (8,9) and, consequently pathogen shedding. Recent work has hypothesized that the dietary balance between sugar, protein and lipid consumption in fruit bats (*Pteropus* spp.) could affect Hendra spillover events in Australia (10). During unfavorable climatic conditions, which negatively affect the flowering of the native eucalyptus trees, bats shift their diet from these native flowers, nectar and pollen rich in sugar and protein, to fruits of introduced plants, that are low in protein and can be high in fat (11–15). This shift to alternative plants often occurs in agricultural areas, where multiple spillover events of Hendra virus from bats to humans *via* horses, a bridging host, have been reported (10,16,17). Potentially, there are several pathways by which diet can affect infection in bats and these can include modifications of the gut microbiome (18,19) or changes in the strength and type of the immune response (18,20–22). However, how nutrition can alter pathogen shedding and, in turn, the likelihood of spillover is poorly understood.

Bats exhibit an immune response that can limit inflammatory responses and facilitate tolerance during viral infections, a pattern that contrasts with the typical mammalian reaction (23,24). However, several stressors can alter the antiviral response and potentially viral shedding (25). For example, bats stressed by anthropogenic disturbances or physiologically challenged, such as during pregnancy or lactation or under poor nutrition, appear to be less capable of controlling infections and conceivably could contribute to higher shedding and transmission (26). These physiological and energetic stressors can alter the processes that allow bats to adapt to energy demands, causing an allostatic overload that suppress the antiviral immune response and increases susceptibility to viral infections (27).

Shifts in diet quality and composition can also alter bat foraging behavior. Studies on black flying foxes (*Pteropus alecto*) in suburban areas of Australia seem to support the general assumption of a negative relationship between food quality and frequency of visits, with more visits to foraging sites during food shortages (11). In general, we could assume that in the absence of food shortage and with easy access to the requited dietary source, bats should be expected to be in good body condition and probably making less frequent and/or shorter visits to the foraging sites. Conversely, under an unbalanced or disrupted diet, bats may make more and/or longer visits to the available foraging sites, which might not completely satisfy their nutritional requirements leading to sub-optimal body conditions.

Here, we explore the dietary-mediated effect of the bat immune response to viral shedding using available laboratory experiments on Jamaican fruit bats (*Artibeus jamaicensis*) maintained at different dietary regimes and infected with the bat-specific influenza A strain H18N11 (28). We develop a mechanistic within-host model, fitted to these data, and test alternative hypotheses on the relationship between diet, immune response and shedding. The selected model is then used to investigate how viral shedding and food consumption, this latter quantity used as a proxy of bat visits to food resources, could provide a working hypothesis on the hazard of spillover events in natural settings. Our findings offer a mechanistic understanding of the role that diet may play for emerging pathogens transmitted by bats and more broadly for zoonotic infections whose dynamics are affected by the dietary requirements of the reservoir hosts.

## Materials and methods

### The system

We used data available derived from laboratory experiments that are outlined by Falvo et al. (28). Briefly, three groups of male Jamaican fruit bats (*Artibeus jamaicensis*) were maintained at different dietary regimes: *i*) an optimal diet of fruits and protein, *ii*) a sub-optimal fat diet of fruits and coconut oil, and *iii*) a sub-optimal diet limited to fruits. After acclimation, bats were inoculated with the bat small-intestine influenza A virus H18N11 (5x10^5^ TCID_50_), and longitudinally monitored for 20 days post infection (DPI). Samples were individually collected to quantify daily viral shedding and, less frequently, immune and metabolic variables (Figure 1). Immune variables and viral load were quantified by RT-PCR, while metabolites were measured using a Sciex 5500 QTRAP mass spectrometer (28). Based on the comprehensive experimental results (28), we developed models using viral RNA from rectal swabs, as a proxy of influenza shedding and two immune variables indicative of the response to the infection/shedding, namely Myxovirus resistance protein 1 (MX1) and Tumor Necrosis Factor Alpha (TNFα). We also used the metabolic marker of intestinal functionality: citrulline, all three variables were quantified from rectal swabs (28). MX1 is a interferon-inducible protein involved in the defense against influenza A viruses (29), while the cytokine TNFα is pivotal in the regulation of the inflammatory response and contributes to the antiviral response (24,30,31). Citrulline is an amino acid involved in the urea cycle and is associated with the improvement of gut immunity, including the integrity of epithelial cells through complex metabolic pathways (32,33). Full details on experimental design, sample collection and processing are reported in Falvo et al. (28).

**Figure 1.**
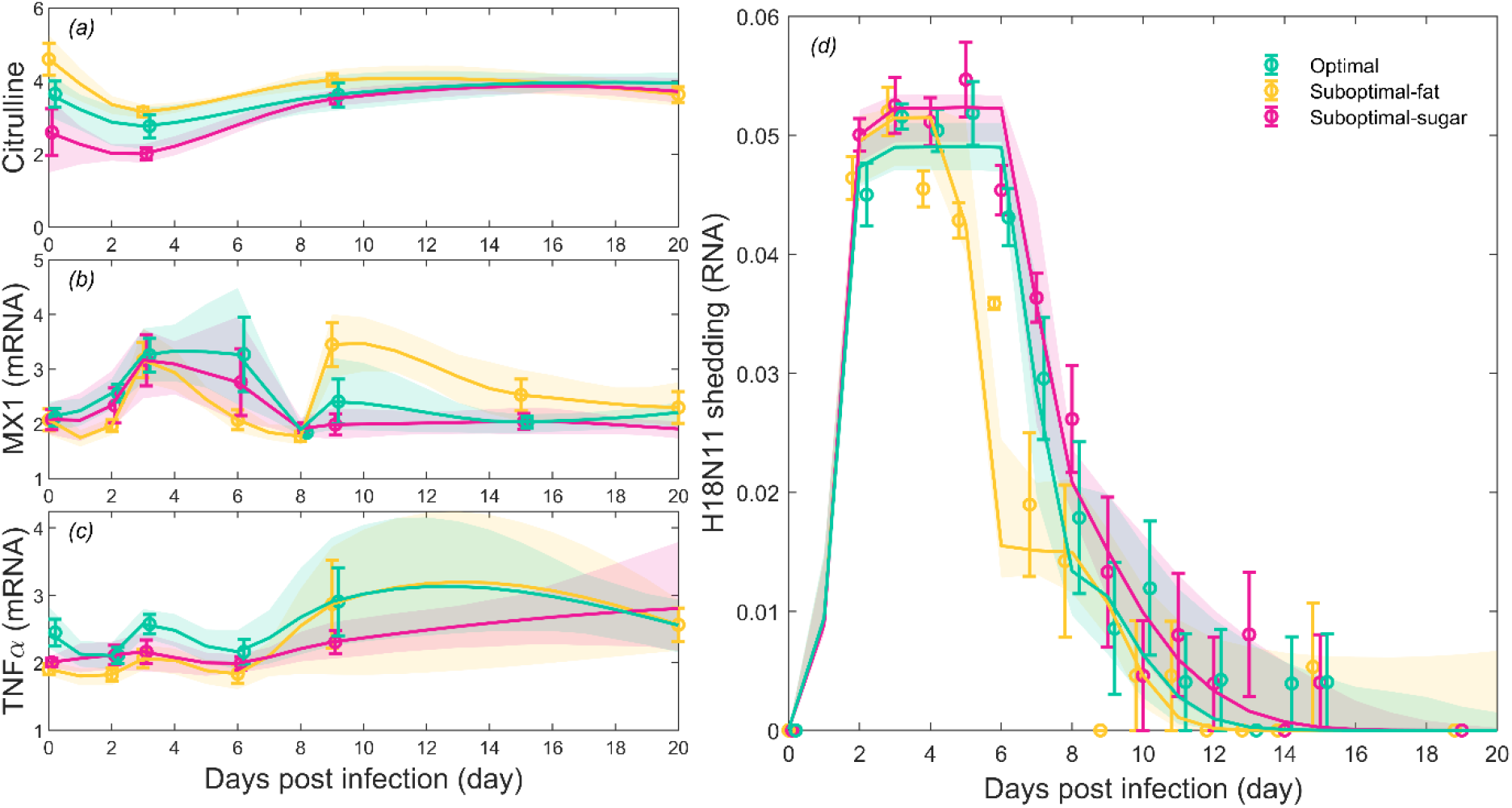
Dietary effect on viral shedding, metabolic and immune variables. (*a*) Citrulline, (*b*) MX1, (*c*) TNFα as potential model covariates and resulting (*d*) viral shedding by time for the optimal (green), sub-optimal fat (yellow) and sub-optimal sugar (purple) bat diet. Points: observed mean +/- S.E; Lines: (*a-c*) interpolated covariates by time, (*b*) simulations from the selected model. Shaded areas: 90% C.I. of model simulations.

### The model

We investigate if the variables MX1, TNFα and citrulline could affect the dynamics of viral shedding, where shedding is assumed to mirror the kinetic of infection in the host’s gut. Five fundamental hypotheses are tested: *HP1:* Viral shedding is controlled by MX1; *HP2:* Viral shedding is controlled by TNFα; *HP3:* Viral shedding is controlled by citrulline; *HP4:* Viral shedding is controlled by citrulline-mediated MX1, *HP5:* Viral shedding is controlled by citrulline-mediated TNFα. These hypotheses are examined with either diet-independent or diet-dependent parameters and with or without a possible delay in the control of shedding, overall generating 19 competing candidate models (Table S1). Given the high number of possible combinations, we limited our analysis to merging those hypothesis options that showed a good performance when tested individually (3 combinations) for a total 22 models tested (Table S1). Below, we present the most complex model that describes the temporal dynamic of viral shedding (*S*) in relation to MX1 (*M*), TNFα (*T*) and citrulline (C), which are assumed to be diet-dependent, *i*=*o* (optimal) or *f* (sub-optimal fat) or *s* (sub-optimal sugar), and characterized by the following differential equation:

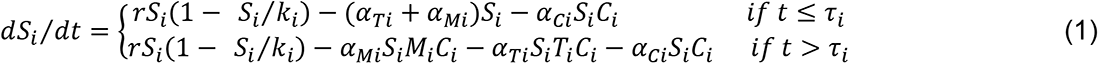

where *t* represents time in days, *r* is the logistic growth rate of viral shedding, *k* is the maximum viral load and *α*_*M*_, *α*_*T*_ and *α*_*C*_ are the neutralization rates of viral shedding induced by MX1, TNFα and citrulline, respectively. The expressions *α*_*Mi*_*M*_*i*_*C*_*i*_, *α*_*Ti*_*T*_*i*_*C*_*i*_ and *α*_*Ci*_*C*_*i*_ describe the immune attack rates, defined as the strength of the viral control by the interaction of immune variables and metabolites. The parameters *τ*_*M*_ and *τ*_*T*_ represent the time delays in the efficacy of MX1 and TNFα, respectively, to control shedding. Parameters with subscript *i* (*k*_*i*_, *α*_*Mi*_, *α*_*Ti*_, and *α*_*Ci*_) identify diet-dependent conditions. In absence of control, viral shedding is assumed to grow logistically to the maximum load (*HP0*, Table S1).

To obtain daily values for the immune and metabolic variables, we performed a Modified Akima piecewise cubic Hermite (Makima) interpolation using the average of each variable for the three bat groups, independently (34,35). The model is then fitted to individual bats from each diet group and parameter values are calibrated by minimizing the following error function, *ERR*, calculated as a weighted sum of viral shedding errors for every individual bat from each diet, using their sample size, *n_o_*, *n_f_*, and *n_s_*, as weights:

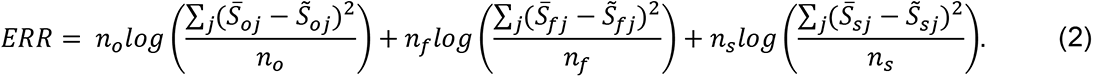

Each error component represents the logarithmic square ratio between observed *S̄_j_* and estimated *S̃_j_* viral shedding values for the three diet groups. The empirical probability distributions of the parameters, and the resulting 90% confidence intervals of their estimates, are calculated via a bootstrap technique (36). The described error function is minimized for each tested model and is equal to the opposite of the double of the log-likelihood (37). Among the calibrated models, we select the best compromise between the goodness of the fit and model parsimony, according to the Akaike Information Criterion (AIC) (38). For each candidate model, the *AIC* score is computed as *AIC = ERR + 2h*, where *h* represents the candidate model complexity, i.e. the number of parameters to calibrate.

### Spillover scenarios

Shedding estimates from the selected model and experimental data on bat food consumption are then used to examine hypothetical spillover scenarios from bat reservoirs to accidental hosts exposed to shedding events. Data on the average food consumed from cage feeders over time (g food/day) were available for the three diets and were normalized by bat body mass (g) (28). We can broadly assume that this measurement is a proxy of bat feeding behavior where the degree of food consumed in a day could be interpreted as the accumulated frequency of visits to and/or time spent foraging at a potential spillover site. The greater the quantity of food taken, the higher is the total frequency of visits and/or the time spent foraging per visit and, thus, the greater is the accumulated pathogen shed at the site. We are aware that bat feeding behavior can have many different patterns which are regulated both by bat physiology and environmental drivers (39), yet, our aim is to present a parsimonious hypothesis for the coarse representation of the shedding-spillover relationship. Importantly, whether this feeding behavior is addressed as frequency or duration of visits, here we are interested in the accumulated daily behavior and related total shedding, under different diets.

Based on the patterns of shedding, we can describe three spillover scenarios, for each diet *i* : *i*) potential spillover (*S_potential,i_*), calculated as the cumulated viral shed over time under no immune control, namely, shedding follows a logistic growth to a maximum load; *ii*) actual spillover (*Sactual,i*), defined as the cumulated viral shed over time under immune control, identified from the selected model framework; and *iii*) effective spillover (*S_effective,i_*), estimated as at point *ii* and corrected by food consumption (*S_t,i_*F_t,i_*), to account for bat’s behavior described either as total frequency or duration of visits. The difference between the first two measurements of viral shedding (*S_potential,i_* - *S_actual,i_*) represents the virus shed neutralized by the immune response while the difference between the last two (*S_actual,i_* - *S_effective,i_*) identifies the virus shed that fails to spillover as outside the time of exposure to an accidental host.

Finally, we perform a sensitivity analysis varying *i*) bat susceptibility (i.e. viral shedding maximum load *k_i_*), *ii)* metabolic-immune response (i.e. mean immune attack rate *α_i_T_i_C_i_*), and *iii*) bat feeding behavior (i.e. daily mean food consumption) and estimate their effects on the hazard of effective viral shedding.

## Results

### Diet affects the metabolic-immune control of viral shedding

An initial examination of the laboratory data shows that the level of H18N11 RNA shedding peaks between day three and six post infection and is controlled within two weeks for each of the three diets (Figure 1d). The peak in shedding is associated with a minimum level of citrulline and an increase of MX1 and TNFα (Figure 1a-c). Both viral shedding and citrulline are significantly affected by the diet regimes (*p*<0.05, Table S2). Bats fed on the sub-optimal fat diet show the fastest control of viral shedding and the highest citrulline level, which contrast with the highest and longest shedding, and lowest citrulline, for the bats on sub-optimal sugar diet (Figure 1); significant statistical differences in the molecular and shedding data between diets support these results (*p*<0.05, Figure S1).

Among the 22 hypothesis-driven models that we tested, the parsimonious model that best captures the temporal control of viral shedding is represented by the synergistic effect of citrulline and TNFα (model M19b) (Figure 1, Tables S1 and S3). This metabolic-immune regulation is diet-dependent: bats from different diets do vary in the strength of this response and their ability to reduce and clear shedding, confirming the experimental observations.

Moreover, this process appears to occur with a time delay, *τ*_*i*_, where the ability to activate an effective immune-metabolic constraint changes with the diet regime. The performance of the selected model slightly underestimates the simulated viral shedding towards the end of the experiment when some bats are still shedding virus, albeit at very low levels (Figure 1d). All the remaining models, including the more complex frameworks, exhibit inferior performances (*ΔAIC*≥6) and are discussed no further (Table S3).

A careful investigation of the selected model shows that the maximum shedding load, which is the maximum amount of virus shed in the absence of host control, is found to be higher in bats fed with the sub-optimal diets when compared to bats from the optimal regime (*k_s_*>*k_f_*>*k_o_*) (Figure 2, Table S4). However, the same sub-optional groups show a stronger control of shedding (i.e. viral mortality rate) induced by the citrulline -TNFα synergistic effect (*α_Ts_*>*α_Tf_*>*α_To_*, Figure 2a,b, Table S4). This control is only effective at 5-7 DPI and the delay is significantly shorter in bats from the sub-optimal fat diet when compared to bats from the other two regimes, which have similar delays (*τ_f_*<*τ_s_*∼*τ_o_*, Figure 2c, Table S4). The temporal kinetic of the immune attack rate, i.e. the product of virus mortality and citrulline-TNFα control, which is indicative of the efficacy to restrain viral shedding, is overall stronger for bats from the sub-optimal fat diet, characterized by faster activation and higher responses (Figure 2d). Bats from the optimal diet show an intermediate ability to control and terminate shedding while the sugar group has an attack rate that is still growing and trying to control shedding at 20 DPI.

**Figure 2.**
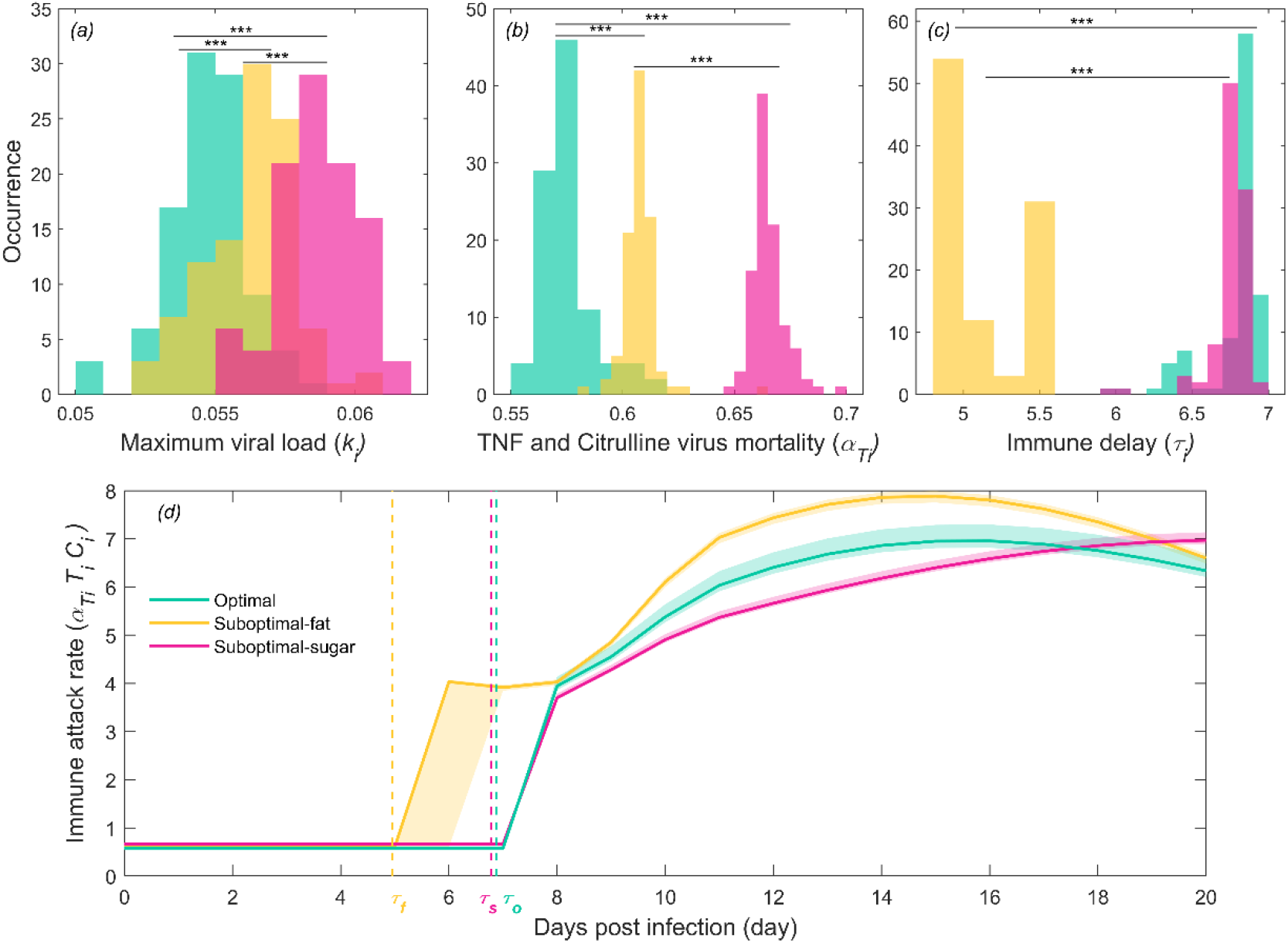
Dietary differences in parameter distributions and immune attack rates. (*a*) Maximum viral shedding load *k_i_*, (*b*) viral-induced mortality rate from Citrulline and TNFα *α_Ti_*, and (*c*) time delay in the immune response *τ_i_*. Between diet groups t-test comparisons: *0.01<*p*<0.05, **0.001<*p*≤0.01, \*\*\**p*≤0.001. (*d*) Immune attack rates, related delays *τ_i_* (vertical dashed lines) and 90% C.I. model simulations (shaded areas).

### Immunity and food consumption affect the hazard of viral spillover

The selected model framework is then used to examine the dietary consequences on the hazard of spillover events, in scenarios where accidental hosts are exposed to viral shedding. In the absence of immune control, bats from the sub-optimal diets shed potentially higher viral loads when compared to bats from the optimal diet (Figure 3a), because of their higher maximum viral load (Table S4). However, under a metabolic-immune control the actual viral shedding is drastically reduced in all three groups, when compared to the potential load (Figure 1d, 3b). Specifically, the metabolic-immune control of bats from the sub-optimal fat diet can block 76% of the potential shedding, this decreased to 70% for the bats on the optimal diet and to 68% for the bats on sub-optimal sugar diet (Figure 3b). We then quantified the effective vial shedding, which is based on the actual shedding corrected by the intake of food from the cage feeders. A preliminary analysis shows that bats from the optimal diet significantly consume the smallest daily quantity of food but have a better return as they maintain the highest body mass. In contrast, bats from the sub-optimal fat diet intake the largest daily amount but have the lowest body mass (Figure S1b,c). Following our rationale, this activity can be interpreted as low daily foraging by the former group and high daily foraging by the latter group, whether this is interpreted as number and/or duration of visits. Based on this, the resulting effective viral shedding is further reduced by bat feeding behavior (Figure 3c). We estimated a 77% reduction from the actual shedding for bats in the optimal diet, whereas this is only reduced to 34%-41% for the bats in the sub-optimal diets. Overall, although bats in the sub-optimal fat diet seem more capable of controlling viral shedding and thus the hazard of spillover, it is the bats from the optimal diet that ultimately are the most successful in reducing such hazard.

**Figure 3.**
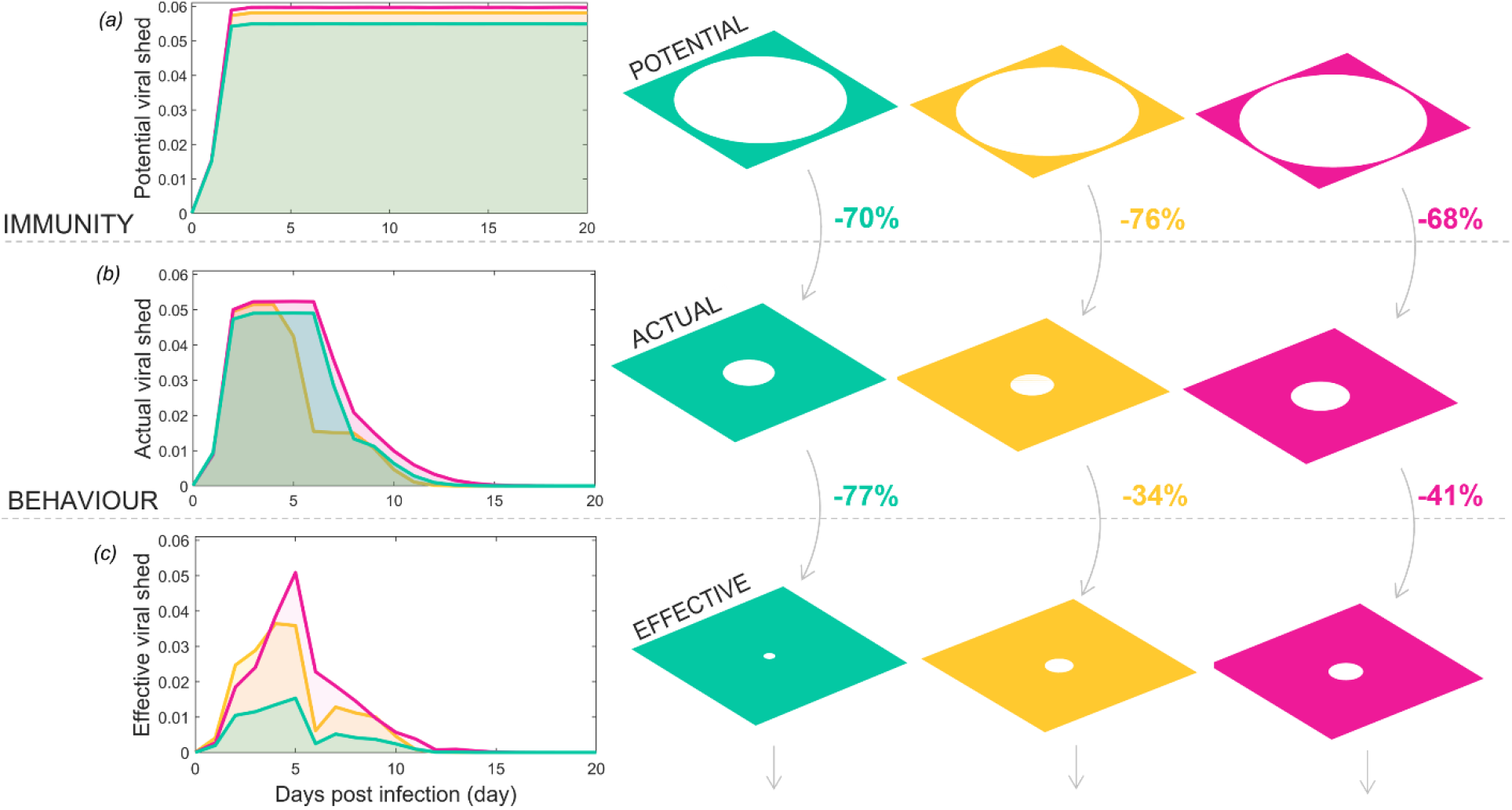
Dietary effects on the barriers to spillover and relative spillover hazard. (*a*) Potential viral shedding, (*b*) actual viral shedding and (*c*) effective viral shedding. The white circle radiuses are proportional to the respective potential, actual, and effective viral shedding accumulated over the infection period (20 days). Percentage of viral shedding blocked by immunity and feeding behavioral barriers, respectively, is reported.

This general outcome is confirmed by the sensitivity analysis that shows that bats on sub- optimal diets have higher capacity to shed virus and consume food but their ability to control shedding can vary, it is either stronger for the fat-diet group or weaker for the sugar-diet group, when compared to bats on optimal diet (Figure S2). Altogether, bats on the optimal diet do relatively better in limiting the possible impact of spillover events.

## Discussion

The goal of this study is to deepen the mechanistic understanding of the dietary impacts on bat immunity, viral shedding dynamics and the hazard of viral spillover. Here, we developed a mechanistic within-host model of viral shedding, fitted to experimental data, to provide a parsimonious explanation of the metabolic-immune control of bats to influenza H18N11 strain under different dietary regimes. This framework was then used to examine hypothetical scenarios of the hazard of pathogen spillover. We showed that diet impacts the onset and strength of the metabolic-immune response leading to contrasting profiles of viral shedding.

When we explored the combined effect of viral shedding and foraging behavior, diet was also important in generating differences in the hazard of infection by an accidental host.

Our selected model indicates that the synergy between the metabolite citrulline and the cytokine TNFα can control and terminate viral shedding. This synergistic response probably involves more complex interactions between components of gut health and the immune system, described in more details in the experimental findings from this system (28). Citrulline activity is associated with other amino acids, ornithine and arginine (28), which were previously found to be linked to protective immune functions involved in the homeostatic maintenance of barrier functions and facilitating the activation of immune cells via endothelial- and inducible-nitric oxide synthase, respectively (40–42). Deficiencies of these metabolites are related to malfunctioning of the inflammatory immune response and the development of conditions like sepsis, bacteremia and endotoxemia in humans (43,44). While these responses need to be tested experimentally in bats, we show that diet-driven differences in citrulline and TNFα levels contribute to affect the baseline inflammatory state of bats pre-infection as well as the response to post-infection, and the consequences on the dynamics of shedding. Bats on the optimal diet presented the highest TNFα values and intermediate levels of citrulline, suggesting that the ability to sustain high inflammation is probably compensated by citrulline, and possibly other amino acids (24) that allow gut good protection against damage. The important role of amino acids in regulating key immune-metabolic pathways and the benefits of arginine/citrulline diet supplementations have been proposed as a novel approach for health and welfare in challenging systems such as farmed fisheries (45,46) and livestock (47,48).

Our modeling results suggest that the overall control of viral shedding by the metabolic-immune response is activated faster and is stronger for the bats on the suboptimal-fat diet. It is possible that the high levels of citrulline and a low TNFα, during the first week post infection, contribute to a faster and more effective metabolic-immune response. The prompt and effective reaction of bats on a fat diet contrasts with the reduced gut health, together with enhanced inflammation and intestinal permeability, found in mice and humans exposed to a fat diet (49,50). While the fundamental mechanism causing these differences is still unclear, it is likely that different metabolic pathways are acting in bats.

No data were available on the level of viral infection in the gastrointestinal tract, this is something that should be considered in the future and important when modeling more precisely the relationship between shedding and immune-metabolic responses. Data on the intensity of infection can also be used for the development of more complex frameworks that explore the molecular mechanisms of inflammation and tissue protection at the local level of the gastrointestinal tissue. This can contribute to explaining why bats can sustain high levels of inflammation. Indeed, the identified parsimonious mechanisms of control of influenza shedding hides a complex chain of processes that are likely involved and that we omitted but that require to be understood and quantified in the laboratory (24,32).

We provide a general working hypothesis of the hazard of pathogen spillover given shedding events and animal food consumption. We propose that bats under dietary shifts from their optimal diet can impact the hazard of viral spillover because of changes in their feeding behavior, whether this is frequency or duration of visits. Foraging behavior is probably affected by body conditions in terms of general health and mass of the individual bats. Indeed, although bats from the sub-optimal diets consume a significantly greater amount of food, they are in poorer body conditions (lower body mass) probably because of their protein-deficient diet which does not fully satisfy their nutritional requirements. Analyses of fecal samples from the field, confirm that the overall protein content of bat diet drives food intake of Pteropodids, which may be forced to over ingest food beyond their energy demand in order to meet their protein requirements (51). In natural conditions, we might also expect these energy unbalances to impact flying activities. For example, *Pteropus alecto* bats were observed to visit more frequently foraging sites during food shortages, compared to conditions of resource abundance (11). Similarly, bat flying activities were found to be significantly higher around fragmented landscape compared to continuous habitats (52). These studies indicate that foraging bats might adjust their movements to density and availability of resources across the landscape (52,53). Moreover, these examples stress the impacts of food availability on bat behavior and energy expenses to seek resources in disrupted landscapes by anthropogenic activities. Our assumption of an association between the quality and quantity of food consumed and the hazard of spillover are drawn using laboratory *A. jamaicensis* as a model system and ultimately need to be confirmed in the field for *Pteropus* spp. Furthermore, there are several other barriers to spillover, in addition to immunity and behavior of the reservoir hosts, that need to be considered when evaluating the dietary effects on spillover hazard (1).

Insights generated from our work can provide testable hypotheses for future work on the bat infection-shedding relationship and the role of immunity to infection, including how individual metabolism and composition of gut microbiome contribute to these patterns. This is critical to gain fundamental knowledge on the ecological mechanisms that contribute to spillover events. Several recent zoonotic outbreaks have been linked to bat-borne viruses (5) and a few of these outbreaks have been suggested to be caused by bat nutritional alterations driven by land use changes (16). Disentangling the link between bat nutrition and ecosystem functioning can inform a proactive response to prevent pathogen spillover with ecological countermeasures aimed at reducing infection and viral shedding at the reservoir source (27,54,55).

## Acknowledgements

This study is funded by NSF-RoL 2133763 grant. The authors are grateful to Dr. Alison Peel for her comments and suggestions to finalize the paper.

