## Supplementary material for "Modeling the dietary effects on bat viral shedding and potential consequences for pathogen spillover": SI

### Supplementary Materials

**Table S1.** Tested models with respective mechanistic hypotheses, viral shedding equations and parameters to be calibrated.

| Model | HP | Viral shedding equation | Parameters to calibrate |
|---|---|---|---|
| M1 | *HP0* | $dS_i/dt = rS_i(1 - S_i/k_i)$ | $r, k_1, k_2, k_3$ |
| M2 | *HP1* | $dS_i/dt = rS_i(1 - S_i/k_i) - \alpha_M S_i M_i$ | $r, k_1, k_2, k_3, \alpha_M$ |
| M3 | *HP1* | $dS_i/dt = rS_i(1 - S_i/k_i) - \alpha_{Mi} S_i M_i$ | $r, k_1, k_2, k_3, \alpha_{M1}, \alpha_{M2}, \alpha_{M3}$ |
| M4 | *HP1* | $dS_i/dt = \begin{cases} rS_i(1 - S_i/k_i) - \alpha_M S_i & if\ t \leq \tau \\ rS_i(1 - S_i/k_i) - \alpha_M S_i M_i & if\ t > \tau \end{cases}$ | $r, k_1, k_2, k_3, \alpha_M, \tau$ |
| M5 | *HP1* | $dS_i/dt = \begin{cases} rS_i(1 - S_i/k_i) - \alpha_M S_i & if\ t \leq \tau \\ rS_i(1 - S_i/k_i) - \alpha_{Mi} S_i M_i & if\ t > \tau \end{cases}$ | $r, k_1, k_2, k_3, \alpha_{M1}, \alpha_{M2}, \alpha_{M3}, \tau$ |
| M4b | *HP1* | $dS_i/dt = \begin{cases} rS_i(1 - S_i/k_i) - \alpha_M S_i & if\ t \leq \tau_i \\ rS_i(1 - S_i/k_i) - \alpha_M S_i M_i & if\ t > \tau_i \end{cases}$ | $r, k_1, k_2, k_3, \alpha_M, \tau$ |
| M5b | *HP1* | $dS_i/dt = \begin{cases} rS_i(1 - S_i/k_i) - \alpha_M S_i & if\ t \leq \tau_i \\ rS_i(1 - S_i/k_i) - \alpha_{Mi} S_i M_i & if\ t > \tau_i \end{cases}$ | $r, k_1, k_2, k_3, \alpha_{M1}, \alpha_{M2}, \alpha_{M3}, \tau_{1,}, \tau_2, \tau_3$ |
| M6 | *HP2* | $dS_i/dt = rS_i(1 - S_i/k_i) - \alpha_T S_i T_i$ | $r, k_1, k_2, k_3, \alpha_T$ |
| M7 | *HP2* | $dS_i/dt = rS_i(1 - S_i/k_i) - \alpha_{Ti} S_i T_i$ | $r, k_1, k_2, k_3, \alpha_{T1}, \alpha_{T2}, \alpha_{T3}$ |
| M8 | *HP2* | $dS_i/dt = \begin{cases} rS_i(1 - S_i/k_i) - \alpha_T S_i & if\ t \leq \tau \\ rS_i(1 - S_i/k_i) - \alpha_T S_i T_i & if\ t > \tau \end{cases}$ | $r, k_1, k_2, k_3, \alpha_T, \tau$ |
| M9 | *HP2* | $dS_i/dt = \begin{cases} rS_i(1 - S_i/k_i) - \alpha_T S_i & if\ t \leq \tau \\ rS_i(1 - S_i/k_i) - \alpha_{Ti} S_i T_i & if\ t > \tau \end{cases}$ | $r, k_1, k_2, k_3, \alpha_{T1}, \alpha_{T2}, \alpha_{T3}, \tau$ |
| M8b | *HP2* | $dS_i/dt = \begin{cases} rS_i(1 - S_i/k_i) - \alpha_T S_i & if\ t \leq \tau_i \\ rS_i(1 - S_i/k_i) - \alpha_T S_i T_i & if\ t > \tau_i \end{cases}$ | $r, k_1, k_2, k_3, \alpha_T, \tau_{1,}, \tau_2, \tau_3$ |
| M9b | *HP2* | $dS_i/dt = \begin{cases} rS_i(1 - S_i/k_i) - \alpha_T S_i & if\ t \leq \tau_i \\ rS_i(1 - S_i/k_i) - \alpha_{Ti} S_i T_i & if\ t > \tau_i \end{cases}$ | $r, k_1, k_2, k_3, \alpha_{T1}, \alpha_{T2}, \alpha_{T3}, \tau_{1,}, \tau_2, \tau_3$ |
| M10 | *HP3* | $dS_i/dt = rS_i(1 - S_i/k_i) - \alpha_C S_i C_i$ | $r, k_1, k_2, k_3, \alpha_C$ |
| M11 | *HP3* | $dS_i/dt = rS_i(1 - S_i/k_i) - \alpha_{Ci} S_i C_i$ | $r, k_1, k_2, k_3, \alpha_{C1}, \alpha_{C2}, \alpha_{C3}$ |
| M12 | *HP3* | $dS_i/dt = rS_i(1 - S_i/k_i) - \alpha_M S_i M_i C_i$ | $r, k_1, k_2, k_3, \alpha_M$ |
| M13 | *HP3* | $dS_i/dt = rS_i(1 - S_i/k_i) - \alpha_{Mi} S_i M_i C_i$ | $r, k_1, k_2, k_3, \alpha_{M1}, \alpha_{M2}, \alpha_{M3}$ |
| M14 | *HP4* | $dS_i/dt = \begin{cases} rS_i(1 - S_i/k_i) - \alpha_M S_i & if\ t \leq \tau \\ rS_i(1 - S_i/k_i) - \alpha_M S_i M_i C_i & if\ t > \tau \end{cases}$ | $r, k_1, k_2, k_3, \alpha_M, \tau$ |
| M15 | *HP4* | $dS_i/dt = \begin{cases} rS_i(1 - S_i/k_i) - \alpha_M S_i & if\ t \leq \tau \\ rS_i(1 - S_i/k_i) - \alpha_{Mi} S_i M_i C_i & if\ t > \tau \end{cases}$ | $r, k_1, k_2, k_3, \alpha_{M1}, \alpha_{M2}, \alpha_{M3}, \tau$ |
| M14a | *HP4* | $dS_i/dt = \begin{cases} rS_i(1 - S_i/k_i) - \alpha_M S_i & if\ t \leq \tau_i \\ rS_i(1 - S_i/k_i) - \alpha_M S_i M_i C_i & if\ t > \tau_i \end{cases}$ | $r, k_1, k_2, k_3, \alpha_M, \tau_{1,}, \tau_2, \tau_3$ |
| M15b | *HP4* | $dS_i/dt = \begin{cases} rS_i(1 - S_i/k_i) - \alpha_M S_i & if\ t \leq \tau_i \\ rS_i(1 - S_i/k_i) - \alpha_{Mi} S_i M_i C_i & if\ t > \tau_i \end{cases}$ | $r, k_1, k_2, k_3, \alpha_{M1}, \alpha_{M2}, \alpha_{M3}, \tau_{1,}, \tau_2, \tau_3$ |
| M16 | *HP5* | $dS_i/dt = rS_i(1 - S_i/k_i) - \alpha_T S_i T_i C_i$ | $r, k_1, k_2, k_3, \alpha_T$ |
| M17 | *HP5* | $dS_i/dt = rS_i(1 - S_i/k_i) - \alpha_{Ti} S_i T_i C_i$ | $r, k_1, k_2, k_3, \alpha_{T1}, \alpha_{T2}, \alpha_{T3}$ |

| M18 | *HP5* | $dS_i/dt = \begin{cases} rS_i(1- \ S_i/k_i) - \alpha_T S_i & if\ t \leq \tau \\ rS_i(1- \ S_i/k_i) - \alpha_T S_i T_i C_i & if\ t > \tau \end{cases}$ | $r, k_1, k_2, k_3, \alpha_T, \tau$ |
|---|---|---|---|
| M19 | *HP5* | $dS_i/dt = \begin{cases} rS_i(1- \ S_i/k_i) - \alpha_T S_i & if\ t \leq \tau \\ rS_i(1- \ S_i/k_i) - \alpha_{Ti} S_i T_i C_i & if\ t > \tau \end{cases}$ | $r, k_1, k_2, k_3, \alpha_{T1}, \alpha_{T2}, \alpha_{T3}, \tau$ |
| M18b | *HP5* | $dS_i/dt = \begin{cases} rS_i(1- \ S_i/k_i) - \alpha_T S_i & if\ t \leq \tau_i \\ rS_i(1- \ S_i/k_i) - \alpha_T S_i T_i C_i & if\ t > \tau_i \end{cases}$ | $r, k_1, k_2, k_3, \alpha_T, \tau_{1,}, \tau_2, \tau_3$ |
| M19b | *HP5* | $dS_i/dt = \begin{cases} rS_i(1- \ S_i/k_i) - \alpha_T S_i & if\ t \leq \tau_i \\ rS_i(1- \ S_i/k_i) - \alpha_{Ti} S_i T_i C_i & if\ t > \tau_i \end{cases}$ | $r, k_1, k_2, k_3, \alpha_{T1}, \alpha_{T2}, \alpha_{T3}, \tau_{1,}, \tau_2, \tau_3$ |
| Full model | *HP3+ HP4+ HP5+* | $dS_i/dt = \begin{cases} rS_i(1- \ S_i/k_i) - (\alpha_{Ti} + \alpha_{Mi})S_i \ - \alpha_{Ci} S_i C_i & if\ t \leq \tau_i \\ rS_i(1- \ S_i/k_i) - \alpha_M S_i M_i C_i - \alpha_{Ti} S_i T_i C_i \ - \alpha_{Ci} S_i C_i & if\ t > \tau_i \end{cases}$ | |

**Table S2.** ANalysis Of VAriance (ANOVA) of the observed viral shedding, MX1, TNFα and Citrulline by time (continuous), diet (categorical), and their interaction. Sample size ( $n_{o,}\ n_f\ n_s$) for the optimal, sub-optimal fat, sub-optimal sugar diet, respectively, are included.

| | **Sum Sq.** | **F** | **p-value** |
|---|---|---|---|
| **Viral shedding (*S*)** $n_o = n_f = n_s = 105$ | | | |
| diet | 0.00104 | 3.17 | 0.0435 |
| time | 0.09768 | 594.78 | $<10^{-16}$ |
| diet* time | 0.00029 | 0.87 | 0.4181 |
| Error | 0.04829 | - | - |
| **MX1(*M*)** $n_o = 56\ n_f = 48\ n_s = 56$ | | | |
| diet | 1.295 | 0.83 | 0.4364 |
| time | 2.029 | 2.61 | 0.1081 |
| diet* time | 2.144 | 1.38 | 0.2546 |
| Error | 118.82 | - | - |
| **TNFα (*T*)** $n_o = 56\ n_f = 48\ n_s = 56$ | | | |
| diet | 2.4871 | 2.58 | 0.0801 |
| time | 5.2443 | 10.88 | 0.0013 |
| diet* time | 0.7881 | 0.82 | 0.444 |
| Error | 54.9307 | - | - |
| **Citrulline (C*)*** $n_o = 28\ n_f = 24\ n_s = 28$ | | | |
| diet | 16.9713 | 10.62 | 0.0001 |
| time | 3.8794 | 4.86 | 0.0306 |

| | | | |
|---|---|---|---|
| diet* time | 6.8253 | 4.27 | 0.0175 |
| Error | 59.1036 | - | - |

**Table S3.** Model selection results. Model complexity (*h*), minimized error for each bat diet ($ERR_o$ for optimal diet, $ERR_f$ for fat diet and $ERR_s$ for sugar diet), total error (*ERR*), Akaike information Criterion (*AIC*) and difference of AIC compared to the best model ($\Delta AIC$). For model description, see Table S1.

| **Model** | ***h*** | $ERR_o$ | $ERR_f$ | $ERR_s$ | ***ERR*** | ***AIC*** | $\Delta AIC$ |
|---|---|---|---|---|---|---|---|
| M1 | 4 | -792.47 | -798.51 | -695.46 | -2286.4 | -2278.4 | 481.4 |
| M2 | 5 | -792.47 | -798.51 | -695.46 | -2286.4 | -2276.4 | 483.4 |
| M3 | 7 | -792.47 | -798.51 | -695.46 | -2286.4 | -2272.4 | 487.4 |
| M4 | 6 | -924.59 | -937.64 | -852.38 | -2714.6 | -2702.6 | 57.2 |
| M5 | 8 | -927.23 | -938.26 | -855.23 | -2720.7 | -2704.7 | 55.1 |
| M4b | 8 | -926.61 | -937.95 | -855.05 | -2719.6 | -2703.6 | 56.2 |
| M5b | 10 | -928.28 | -938.40 | -855.60 | -2722.3 | -2702.3 | 57.5 |
| M6 | 5 | -833.39 | -857.94 | -762.42 | -2453.8 | -2443.8 | 316 |
| M7 | 7 | -857.42 | -858.17 | -768.17 | -2483.8 | -2469.8 | 290 |
| M8 | 6 | -968.01 | -946.71 | -817.19 | -2731.9 | -2719.9 | 39.9 |
| M9 | 8 | -968.24 | -951.85 | -822.19 | -2742.3 | -2726.3 | 33.5 |
| M8b | 8 | -968.56 | -958.02 | -817.05 | -2743.6 | -2727.6 | 32.2 |
| M9b | 10 | -969.52 | -969.99 | -817.63 | -2757.1 | -2737.1 | 22.7 |
| M10 | 5 | -934.20 | -856.08 | -665.22 | -2455.5 | -2445.5 | 314.3 |
| M11 | 7 | -952.75 | -855.54 | -666.73 | -2475.0 | -2461 | 298.8 |
| M12 | 5 | -792.47 | -798.51 | -695.46 | -2286.4 | -2276.4 | 483.4 |
| M13 | 7 | -792.47 | -798.51 | -695.46 | -2286.4 | -2272.4 | 487.4 |
| M14 | 6 | -923.66 | -937.42 | -848.15 | -2709.2 | -2697.2 | 62.6 |
| M15 | 8 | -923.70 | -937.67 | -848.27 | -2709.6 | -2693.6 | 66.2 |
| M14b | 8 | -923.69 | -937.65 | -848.23 | -2709.6 | -2693.6 | 66.2 |
| M15b | 10 | -923.69 | -937.65 | -848.23 | -2709.6 | -2689.6 | 70.2 |
| M16 | 5 | -859.18 | -852.76 | -739.67 | -2451.6 | -2441.6 | 318.2 |
| M17 | 7 | -922.96 | -834.15 | -737.95 | -2495.1 | -2481.1 | 278.7 |
| M18 | 6 | -930.21 | -895.22 | -817.39 | -2642.8 | -2630.8 | 129 |
| M19 | 8 | -961.79 | -944.47 | -846.38 | -2752.6 | -2736.6 | 23.20 |
| M18b | 8 | -974.76 | -968.21 | -812.38 | -2755.4 | -2739.4 | 20.40 |
| M19b | 10 | -977.97 | -968.63 | -833.22 | -2779.8 | -2759.8 | 0 |

| | | | | | | | |
|---|---|---|---|---|---|---|---|
| M5b+M11 | 13 | -974.44 | -948.74 | -853.78 | -2777.0 | -2751 | 8.80 |
| M5b+M19b | 13 | -977.97 | -968.63 | -833.22 | -2779.8 | -2753.8 | 6 |
| M11+M19b | 14 | -977.97 | -968.63 | -833.22 | -2779.8 | -2751.8 | 8 |
| M5b+M11+M19b | 16 | -977.97 | -968.63 | -833.22 | -2779.8 | -2747.8 | 12 |

**Table S4.** Selected model (M19b) estimated parameter values, unit of measure and 90% C.I.

| **Parameters** | **Value** | **Unit of measure** | **90% CI** |
|---|---|---|---|
| $r$ | 5.3348 | day$^{-1}$ | [5.2197; 5.9593] |
| $k_o$ | 0.0549 | RNA | [0.0526; 0.0569] |
| $k_f$ | 0.0581 | RNA | [0.0535; 0.0587] |
| $k_s$ | 0.0597 | RNA | [0.0559; 0.0608] |
| $\alpha_{To}$ | 0.5734 | (mRNA day)$^{-1}$ | [0.5616; 0.6014] |
| $\alpha_{Tf}$ | 0.6077 | (mRNA day)$^{-1}$ | [0.5973; 0.6158] |
| $\alpha_{Ts}$ | 0.6630 | (mRNA day)$^{-1}$ | [0.6555; 0.6786] |
| $\tau_o$ | 6.8455 | day | [6.3312; 6.9290] |
| $\tau_f$ | 4.9756 | day | [4.9344; 5.4993] |
| $\tau_s$ | 6.8044 | day | [6.5184; 6.8561] |

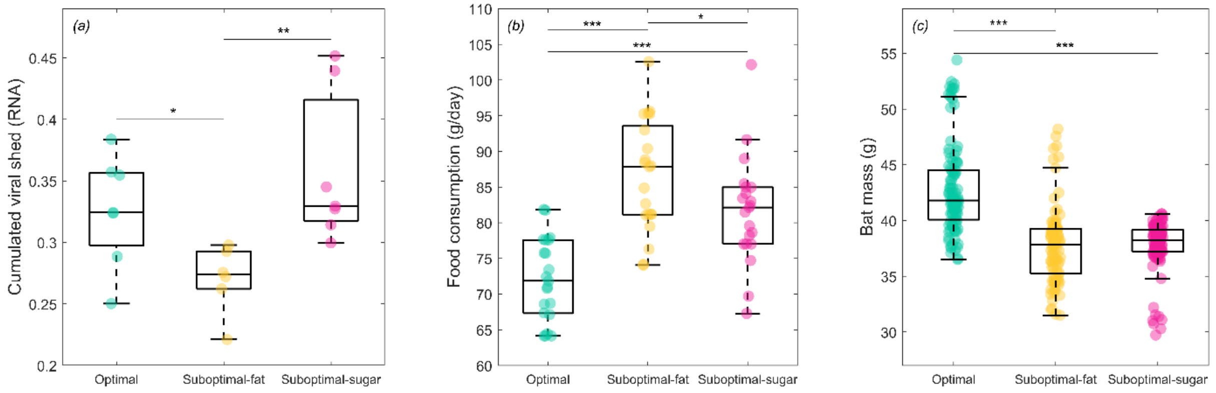


**Figure S1**. Observed bat data by diets. Boxplots of (*a*) observed cumulated viral shed by individual bats, (*b*) average daily food consumption by bat cage and (*c*) bat daily body mass by optimal (green), sub-optimal fat (yellow) and sub-optimal sugar (purple) diet group. Between diet groups t-test comparisons: *0.01<$p$<0.05, **0.001<$p$≤0.01, ***$p$≤0.001.

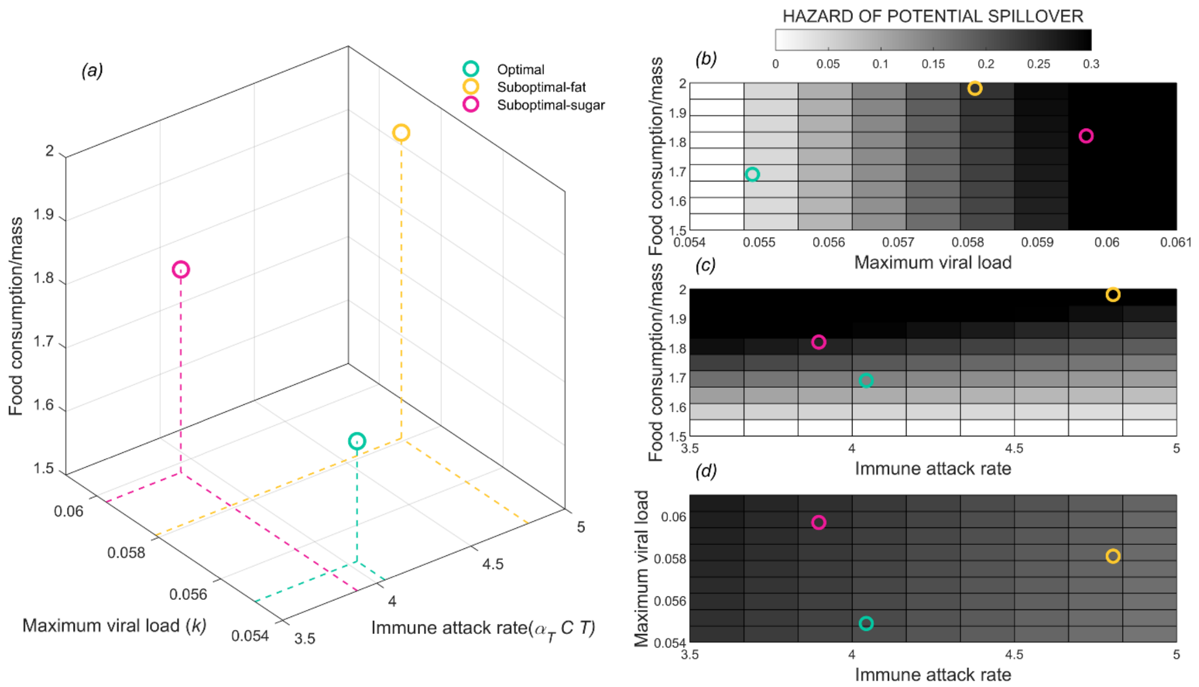


**Figure S2**. Sensitivity analysis. (*a*) Maximum viral load ($k_i$, x-axis), mean immune attack rate ($\alpha_{Ti}T_iC_i$, y-axis) and food consumption/bat mass (z-axis) of bats belonging to the three diet groups. (*b-d*) Hazard of effective viral shedding (gray color scale) by (*b*) viral carrying capacity and food consumption, (*c*) immunity and food consumption, and (*c*) immunity and viral carrying capacity. For each panel, the third variable not represented is kept constant and equal to the mean across diets.
